# Why are tropical mountain passes ‘low’ for some species? Genetic and stable-isotope tests for differentiation, migration, and expansion in elevational generalist songbirds

**DOI:** 10.1101/161380

**Authors:** Chauncey R. Gadek, Seth D. Newsome, Elizabeth J. Beckman, Andrea C. Chavez, Spencer C. Galen, Emil Bautista, Christopher C. Witt

## Abstract

Most tropical bird species have narrow elevational ranges, likely reflecting climatic specialization. This is consistent with Janzen’s Rule, the tendency for mountain passes to be effectively ‘higher’ in the tropics. Hence, those few tropical species that occur across broad elevational gradients raise questions. Are they being sundered by diversifying selection along the gradient? Does elevational movement cause them to resist diversification or specialization? Have they recently expanded, suggesting that elevational generalism is short-lived in geological time? Here we tested for differentiation, movement, and expansion in four elevational generalist songbird species on the Andean west slope. We used morphology and mtDNA to test for genetic differentiation between high- and low-elevation populations. Morphology differed for House Wren (*Troglodytes aedon*) and Hooded Siskin (*Spinus magellanicus*), but not for Cinereous Conebill (*Conirostrum* cinereum) and Rufous-collared Sparrow (*Zonotrichia capensis*), respectively. mtDNA was structured by elevation only in *Z. capensis*. To test for elevational movements, we measured hydrogen isotope (δ^2^H) values of metabolically inert feathers and metabolically active liver. δ^2^H data indicated elevational movements by two tree- and shrub-foraging species with moderate-to-high vagility (*C. cinereum* and *S. magellanicus*), and sedentary behavior by two terrestrial-foraging species with low-to-moderate vagility (*T. aedon* and *Z. capensis*). In *S. magellanicus*, elevational movements and lack of mtDNA structure contrast with striking morphological divergence, suggesting strong diversifying selection on body proportions across the ∼50 km gradient. All species except *C. cinereum* exhibited mtDNA-haplotype variation consistent with recent population expansion across the elevational gradient.In sum,three out of four elevational generalist species underwent genetic divergence despite gene flow, two of four make seasonal elevational movements, and three of four have recently expanded. In different ways, each species defies the tendency for tropical birds to have long-term stable distributions and sedentary habits. We conclude that tropical elevational generalism is rare due to evolutionary instability.

## Introduction

Elevational gradients cause profound eco-climatic variation across short distances. As a result, mountains are important hotspots of biodiversity (e.g. Sanders 2002, Grytnes and Vetaas 2002, McCain 2003) and drivers of diversification (e.g. Ribas et al. 2007, Freeman 2015, Galen et al. 2015, Benham and Witt 2016, Bertrand et al. 2016). As elevation increases, organisms must cope with reduced temperature, humidity, air density, and partial pressure of oxygen (*P*O_2_), and increased exposure to UV radiation. The seasonally stable climatic gradients imposed by tropical mountains have been implicated in global latitudinal biodiversity gradients via Janzen’s Rule (Janzen 1967). Janzen’s Rule holds that mountain passes are effectively ‘higher’ in the tropics because seasonal thermal stability on tropical slopes has led to elevational specialization and discouraged dispersal across elevations. With increased specialization and reduced dispersal, tropical mountains promote allopatric diversification. As a result, tropical species should have narrower elevational ranges than temperate ones. Although empirical support for that prediction is mixed (McCain 2009), narrow elevational distributions are the predominant pattern for tropical montane landbirds, including songbirds (Terborgh 1971, Parker et al. 1996, Jankowski et al. 2013).

The tendency for Andean songbird species to have narrow elevational ranges is strong, as evidenced by their average elevational range breadth of only ∼1250 m on a habitable elevational gradient spanning > 5000 m (Parker et al. 1996). These narrow elevational distributions reflect firmly established elevational limits to species distributions. However, a small number of species defy this pattern, particularly on the west slope of the Andes. Among tropical Andean songbirds, ∼4% of species inhabit elevational ranges broader than 3000 m (Parker et al. 1996) and represent exceptions to Janzen’s Rule because they encounter a broad range of climatic conditions. We offer three non-mutually exclusive explanations for the existence of these broad elevational ranges at tropical latitudes. The first explanation is that genetic fit with the environment is facilitated by cryptic population genetic structure (Milá et al. 2009, 2010); such structure can be either genome-wide or limited to functional loci that may be subject to natural selection that is strong enough to overcome the homogenizing effects of gene flow. A second possible explanation is that individuals make elevational movements to track resources; such movement could prevent specialization by inhibiting spatially variable selection and the isolation of subpopulations along an elevational gradient. A third possible explanation is that population expansion across elevational gradients occurs periodically, but the resulting mismatch between genes and environment leads to subsequent specialization via range contraction or genetic differentiation. Tests of these mechanisms could help explain the rarity of tropical elevational generalism.

Several studies suggest that bird species can diversify along elevational gradients (McCormack and Smith 2008, Cheviron and Brumfield 2009, Milá et al. 2009, McCormack and Berg 2010, Galen et al. 2015). Whether this diversification can lead to speciation without cessation of gene flow is uncertain, but it is theoretically possible (Hua 2016). While hypoxia, cold temperatures, and high UV exposure associated with high elevations are known to cause rapid evolutionary emergence of novel phenotypes (e.g. Beall et al. 2010, Simonson et al. 2010, Galen et al. 2015), homogenizing gene flow between high and low populations is expected to inhibit functional divergence and speciation (Rundle and Nosil 2005). Differentiation with gene flow has been shown under some circumstances (e.g. Kirkpatrick and Barton 1997, Milá et al. 2009, Gutiérrez-Pinto et al. 2014, Benham and Witt 2016), but gene flow generally limits the extent of local adaptation. For example, Benham and Witt (2016) found that the degree of hummingbird bill size differentiation across a climatic gradient was constrained where habitats were contiguous. For sedentary elevational generalists, selection that varies along elevational gradients should lead to differentiation between high- and low-elevation populations at functional loci (Storz and Kelly 2008, Natarajan et al. 2015). The latter process can lead to speciation if functional alleles have pleiotropic effects on reproductive isolation (Hua 2016). Discontiguous habitat along an elevational gradient could facilitate functional and neutral divergence via isolation. Alternatively, elevational movements could directly hinder such divergence.

Elevational migration comprises short-distance movements to track elevation-specific resource pulses that are important for reproduction (Loiselle and Blake 1991, Johnson and Maclean 1994, Boyle 2017). It has been documented in numerous animal species (Hunt et al. 1999, McGuire and Boyle 2013, Voigt et al. 2013), particularly birds (Loiselle and Blake 1991, Chaves-Campos et al. 2003, Hobson et al. 2003, Boyle 2010, Newsome et al. 2015, Villegas et al. 2016). If elevational generalist species undertake seasonal movements, they may be able to track resource pulses or temperature niches (Boyle 2017), but individuals would also experience variable air density, *P*O_2_, and UV radiation that vary predictably with elevation during all seasons (West 1996). These individual movements would reduce the spatial variability of selection and facilitate gene flow that inhibits elevational divergence (Arguedas and Parker 2000). Despite the dramatic elevational gradients of the New World tropics, previous studies in the region have found limited evidence of elevational migration, and most elevational movements that have been documented are small in magnitude (Hobson et al. 2003, Boyle 2010, Hardesty and Fraser 2010, Boyle et al. 2010, Villegas et al. 2016). In contrast, partial or full elevational migration may be more common at temperate and subtropical latitudes in the Andes (e.g., Newsome et al. (2015)). Remarkably, the frequency and extent of elevational migration for most tropical Andean bird species remains unstudied, especially in small-bodied passerine species for which satellite-tracking technologies have yet to be applied.

Hydrogen isotope (δ^2^H) values of bird tissues can be used to characterize latitudinal and elevational movements (Hobson 1999, Bowen et al. 2005). The δ^2^H of precipitation varies predictably with respect to a variety of physicochemical processes (Dansgaard 1964, Estep and Dabrowski 1980, Estep 1981, Rubenstein and Hobson 2004). As water vapor rises on the windward side of a mountain range, it cools and condenses, and water containing the heavier isotope of hydrogen (deuterium) is the first to condense. This produces a systematic relationship between elevation and δ^2^H of local precipitation, resulting in lapse rates of 4 – 8‰ per 100 m (Poage 2001). δ^2^H values of primary producers reflect local precipitation, and consumers integrate δ^2^H values of food and water such that their tissues have δ^2^H that is higher than their food but lower than their water (Estep and Dabrowski 1980, Hobson et al. 1999, Birchall et al. 2005, Wolf et al. 2013). A few studies have utilized the elevational lapse rate in the δ^2^H of precipitation to assess elevational movements (Hobson et al. 2003, Hardesty and Fraser 2010, Newsome et al. 2015). Most δ^2^H-based studies have analyzed feathers, a metabolically inert tissue that records ecological information only during molt, which may only last a few weeks (Hobson et al. 2003, Pérez and Hobson 2007, Wunder 2012, Hobson et al. 2012). More recently, a multi-tissue approach comparing δ^2^H values of metabolically active tissues (i.e. blood, muscle, liver) with metabolically inert feathers offers the potential to reveal utilization of high versus low elevation resources during different periods of the annual life cycle. (Mazerolle and Hobson 2005, Hardesty and Fraser 2010, Newsome et al. 2015, Villegas et al. 2016).

Here we test our explanations for broad elevational distributions in four tropical songbird species: Cinereous Conebill (*Conirostrum. Cinereum),* Hooded Siskin (*Spinus magellanicus*), House Wren (*Troglodytes aedon*), and Rufous-collared Sparrow (*Zonotrichia capensis*). We used morphological and genetic data to test for genetic differentiation and signals of expansion along the gradient. We examined δ^2^H in metabolically inert (feathers) and active (liver) tissues to test for elevational movements in species with varying foraging strata and dispersal abilities. Our analyses suggest elevational movement in *C. cinereum* and *S. magellanicus*, genetic differentiation in *S. magellanicus, T. aedon*, and *Z. capensis*, and recent demographic expansion in all species except *C. cinereum*. These patterns of elevational movement and diversification are consistent, in part, with all three proposed explanations for the relative rarity of tropical elevational generalists.

## Methods

### Morphometric analyses

To test for genetic differentiation across the elevational range at the loci underlying functional morphological traits, we compared sizes of four traits. For four focal species, *C. cinereum, S. magellanicus, T. aedon*, and *Z. capensis*, we compared morphological measurements between populations at high (>3000 m) and low (<1000 m) elevations. We measured culmen, wing chord, tail, and tarsus from museum specimens listed in Appendix 5. We used PCA to visualize the morphometric data, and MANOVA or non-parametric Kruskal-Wallace tests to compare high and low groups.

To help interpret the results of this study, we assessed relative flight capabilities of our four study species. To do so, we compared relative flight muscle mass and hand-wing index, measures that are known to correlate with flight ability and dispersal propensity (Kipp 1959, Dawideit et al. 2009, Burney and Brumfield 2009, Claramunt et al. 2012, Wright et al. 2014, 2016).

### mtDNA population differentiation

To test for mtDNA differentiation across the elevational range, we analyzed published (Cheviron and Brumfield 2009, Galen and Witt 2014) and original mtDNA sequence data from high-elevation (>3000 m) and low-elevation (<1000 m) specimens listed in Appendix 7. We tested for elevational population genetic structure by estimating Fst and Φ st between elevational zones using Arlequin v3.5 (Excoffier and Lischer 2010).

### Stable isotope measurements

To test for elevational movements, we used mass spectrometry to measure δ^2^H from liver, contour feathers, and secondary flight feathers from museum specimens of our four focal taxa that were collected over the last decade on the west slope of the Andes in central Peru (Appendix 4). Technical details are described in Supplementary Material, Appendix 1.

### δ^2^H hypothetical framework

We sought to exploit the ubiquitous trend of decreasing precipitation δ^2^H with increasing elevation (Poage 2001, Gonfiantini et al. 2001) to test for short distance elevational migration (Hobson et al. 2003, Newsome et al. 2015, Villegas et al. 2016). While few precipitation δ^2^H datasets exist for the west slope of the Peruvian Andes (IAEA/WMO 2015), elevational δ^2^H lapse rates for other Andean regions range from 4 to 8‰ per 100m (Niewodniczanski et al. 1981, Rozanski and Araguás-Araguás 1995, Araguás-Araguás et al. 1998, Poage 2001). Our study system on the west slope of the Peruvian Andes is influenced, at low elevations, by ^2^H-enriched fog (Scholl et al. 2010) coupled with sporadic precipitation, and at higher elevations by ^2^H-depleted precipitation resulting from a combination of temperature-dependent fractionation and Rayleigh distillation (Dansgaard 1964, Poage 2001). These disparate isotopic inputs likely produce elevational lapse rates in precipitation δ^2^H (4–8‰ per 100m) that are comparable to those reported by Poage and Chamberlain (2001). Yet, seasonal changes in precipitation δ^2^H that are of equal or greater magnitude than elevational variation in δ^2^H may obscure expected elevational trends in δ^2^H of metabolically active bird tissue (Gonfiantini et al. 2001, IAEA/WMO 2015, Villegas et al. 2016).

To account for seasonal variation in precipitation δ^2^H values across the west slope of the Peruvian Andes, we used precipitation δ^2^H data collected from 2006–2008 in Marcapomacocha, Peru (∼4400m) from the Global Network of Isotopes in Precipitation (GNIP) (IAEA/WMO, 2015). Marcapomacocha, at the top of the transect where most of our specimens were collected (Fig. S1), is the only site on the west slope of the Peruvian Andes for which multiple years of monthly measurements of precipitation δ^2^H values exist. We used monthly mean precipitation δ^2^H values (δ^2^H_month_) obtained from the Marcapomacocha to account for effects of seasonal fluctuations in precipitation δ^2^H in our model of elevational effects on δ^2^H of metabolically active bird liver tissue. The dearth of available data on precipitation δ^2^H at other elevations requires us to assume that seasonal fluctuations in precipitation δ^2^H occur similarly across elevations.

Differences in migratory behavior among elevational-generalist songbird species will be reflected in how their metabolically inert versus active tissues differentially integrate elevational versus seasonal trends in local precipitation δ^2^H. Feather tissues are grown over the course of a few weeks, typically during the dry season (June to August; Fig. S2), after which they become metabolically inert. Thus, δ^2^H_feather_ should not be influenced by the date of sampling, which is captured in our models by δ^2^H_month_.

δ^2^H_feather_ values are expected to vary with elevation of capture in sedentary birds, but not in migratory ones that often will have shifted in elevation between the date of molt and the date of sampling. Conversely, liver tissue is metabolically active so δ^2^H_liver_ should reflect both seasonal (δ^2^H_month_) and elevational variation in precipitation δ^2^H, regardless of whether the bird is sedentary or migratory. Thus, regardless of the date on which a bird was collected, liver δ^2^H values should be predicted in part by seasonal fluctuations in precipitation δ^2^H, which are reflected in our models as δ^2^H_month_.

Elevational movements of individual birds are predicted to influence the relationship between tissue δ^2^H and elevation. If elevational migration occurs, we expect that any correlation between δ^2^H and elevation of capture would be diminished in metabolically inert tissues, and potentially also in metabolically active ones. Because elevational migrant species are less likely to have grown feathers at the elevation of capture, we expect δ^2^H values for inert feathers to lack a trend with elevation of capture.

### δ^2^H statistical analyses

We compared intra- and interspecific δ^2^H_tissue_ values using ANOVA and Kruskal-Wallis or Wilcoxon Rank-Sum tests where applicable. To test for elevational and seasonal effects on δ^2^H we evaluated sets of linear models for each species and tissue type using AIC_c_. For δ^2^H_liver_, we compared models containing all possible combinations of the intercept and three continuous predictor variables: elevation, precipitation δ^2^H_month_ for the sampling date, and latitude. For δ^2^H_feather_ we excluded models that included precipitation δ^2^H_month_ because metabolically inert tissues should be independent of precipitation-δ^2^H at the date of capture. We included latitude as a potentially confounding variable, but we did not consider models with latitude as the sole predictor variable. Furthermore, we excluded models that performed worse by AIC_c_ than a nested version of the same model (Arnold 2010).

### mtDNA test of recent population expansion

To test for recent population expansion, we used DnaSP v5 (Librado and Rozas 2009) to estimate Tajima’s D (Tajima 1996), Fu’s F (Fu 1997), and we used mismatch distributions to evaluate the distribution of pairwise divergence between individuals in a population. Using the mismatch distributions, we calculated the raggedness index (r), with raggedness expected to be elevated under a stable population relative to an expanding one (Harpending 1994). We inferred population expansion when both Tajima’s D and Fu’s F were significantly negative and there was no significant raggedness.

Detailed methods are reported in the Supplementary Material, Appendix 1.

## Results

### Precipitation δ^2^H

GNIP data (IAEA/WMO 2015) revealed striking seasonal variation for the Marcapomacocha GNIP site (Fig. S1). Mean δ2H_month_ was −117±33‰ during the wet season (Oct–May) and −40±19‰ during the dry season (June –Sep). The large (∼100‰) difference between precipitation δ^2^H in the wet season and dry season (Fig. S1) represents a confounding factor that requires careful consideration when attempting to interpret δ^2^H values of metabolically active tissues collected along an elevational gradient.

### Tissue δ^2^H

Comparisons of linear models to explain tissue δ^2^H values are reported in Table 1. Neither latitude nor elevation explained variation in feather-δ^2^H for *C. cinereum* or *S. magellanicus*. In contrast, top models for both feather types of *T. aedon* included elevation of capture as the sole predictor variable, although only δ^2^H_contour_ values were significantly negatively correlated with elevation of collection (*t value*: −2.67, *P* = 0.01; Fig. 1). Similarly, top models for *Z. capensis* feather-δ^2^H included only elevation of capture as a predictor variable, and both δ^2^H_contour_ (*t value*: −2.96, *P* < 0.01) and δ^2^H_secondary_ (*t value*: −3.99, *P* < 0.001) were significantly negatively correlated with elevation of capture (Fig. 1; Table 1).

**Table 1.** Comparison of models to explain δ^2^H_tissus_ values as a function of elevation (elev), seasonal variation in precipitation δ^2^H (δ^2^H_month_), and latitude (lat) for each of the four-focal species. ^2^H_month_ was excluded from comparisons for δ^2^H_feather_ values (dark gray boxes). All combinations of predictor variables were tested against δ^2^H_liver_. Models that scored lower than nested versions of themselves were removed, following Arnold (2010). For each species and tissue type, models with lowest AIC_c_, ΔAIC_c_ of 0, and highest weight are bolded.

| species | model | $\delta^2\text{H}_{\text{contour}}$ | | | $\delta^2\text{H}_{\text{secondary}}$ | | | $\delta^2\text{H}_{\text{liver}}$ | | |
| --- | --- | --- | --- | --- | --- | --- | --- | --- | --- | --- |
| | | AIC<sub>c</sub> | $\Delta\text{AIC}_c$ | weight | AIC<sub>c</sub> | $\Delta\text{AIC}_c$ | weight | AIC<sub>c</sub> | $\Delta\text{AIC}_c$ | weight |
| *C. cinereum* | ~ 1 | **299.83** | **0.00** | **1.00** | **322.68** | **0.00** | **1.00** | 328.89 | 2.24 | 0.25 |
|  | ~ elev |  |  |  |  |  |  |  |  |  |
|  | ~ elev + lat |  |  |  |  |  |  |  |  |  |
| | ~ $\delta^2\text{H}_{\text{month}}$ | | | | | | | **326.66** | **0.00** | **0.75** |
| | ~ elev + $\delta^2\text{H}_{\text{month}}$ | | | | | | | | | |
| | ~ $\delta^2\text{H}_{\text{month}}$ + lat | | | | | | | | | |
| | ~ elev + $\delta^2\text{H}_{\text{month}}$ + lat | | | | | | | | | |
| *S. magellanicus* | ~ 1 | **188.59** | **0.00** | **1.00** | **176.93** | **0.00** | **1.00** | 182.08 | 14.34 | 0.00 |
|  | ~ elev |  |  |  |  |  |  |  |  |  |
|  | ~ elev + lat |  |  |  |  |  |  |  |  |  |
| | ~ $\delta^2\text{H}_{\text{month}}$ | | | | | | | 174.54 | 6.80 | 0.03 |
| | ~ elev + $\delta^2\text{H}_{\text{month}}$ | | | | | | | 172.62 | 4.87 | 0.07 |
| | ~ $\delta^2\text{H}_{\text{month}}$ + lat | | | | | | | 172.05 | 4.30 | 0.09 |
| | ~ elev + $\delta^2\text{H}_{\text{month}}$ + lat | | | | | | | **167.74** | **0.00** | **0.81** |
| *T. aedon* | ~ 1 | 249.30 | 4.76 | 0.08 | 257.72 | 0.90 | 0.39 | 275.11 | 9.15 | 0.00 |
|  | ~ elev | **244.55** | **0.00** | **0.92** | **256.83** | **0.00** | **0.61** |  |  |  |
|  | ~ elev + lat |  |  |  |  |  |  |  |  |  |
| | ~ $\delta^2\text{H}_{\text{month}}$ | | | | | | | 266.23 | 0.27 | 0.31 |
| | ~ elev + $\delta^2\text{H}_{\text{month}}$ | | | | | | | 266.03 | 0.07 | 0.34 |
| | ~ $\delta^2\text{H}_{\text{month}}$ + lat | | | | | | | | | |
| | ~ elev + $\delta^2\text{H}_{\text{month}}$ + lat | | | | | | | **265.95** | **0.00** | **0.35** |
| *Z. capensis* | ~ 1 | 272.86 | 5.72 | 0.05 | 264.56 | 11.11 | 0.00 | 264.71 | 23.80 | 0.00 |
|  | ~ elev | **267.14** | **0.00** | **0.95** | **253.45** | **0.00** | **1.00** | 250.44 | 9.53 | 0.01 |
|  | ~ elev + lat |  |  |  |  |  |  | 248.86 | 7.95 | 0.02 |
| | ~ $\delta^2\text{H}_{\text{month}}$ | | | | | | | 264.35 | 23.44 | 0.00 |
| | ~ elev + $\delta^2\text{H}_{\text{month}}$ | | | | | | | 249.16 | 8.25 | 0.02 |
| | ~ $\delta^2\text{H}_{\text{month}}$ + lat | | | | | | | | | |
| | ~ elev + $\delta^2\text{H}_{\text{month}}$ + lat | | | | | | | **240.91** | **0.00** | **0.96** |

**Figure 1.**
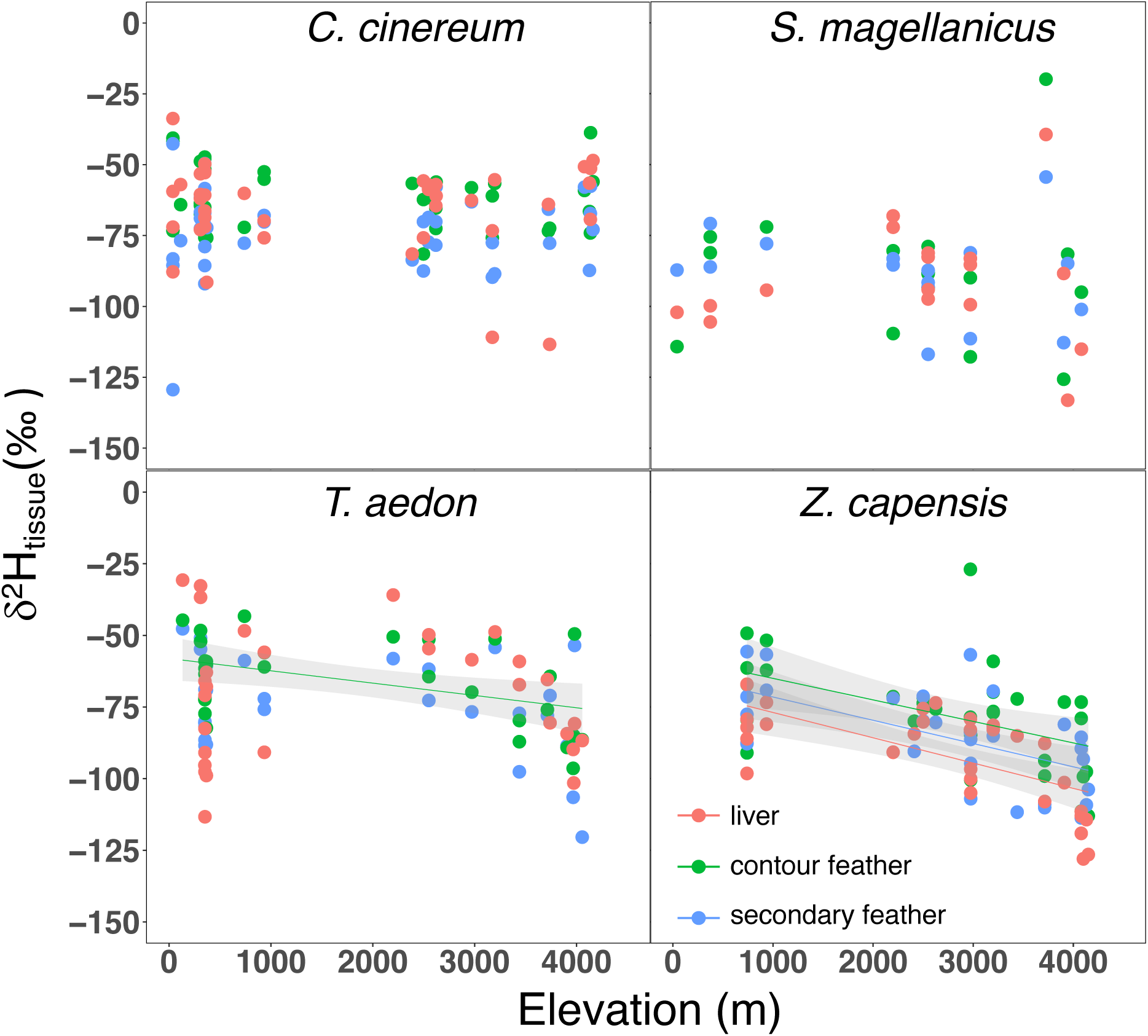
D^2^H values contour feather (green), secondary flight feather (blue), and liver (red) plotted against elevation of capture for *C. cinereum, S. magellanicus, T. aedon*, and *Z. capensis*; sample sizes for each species are reported in Table 2; statistics for linear relationships are provided in Table 3. Best-fit lines are shown for linear regressions that are significant at p < 0.05 and gray bands represent 95% confidence intervals.

δ^2^H_liver_ varied significantly among species (*F value*: 18.59, DF = 3, *P* < 0.001), with values for the two insectivorous species (*C. cinereum* and *T. aedon*) significantly higher than those of the two granivorous species (*S. magellanicus* and *Z. capensis*). For all four species, the best models with δ^2^H_liver_ as a response variable included δ^2^H_month_ a s predictor variable (Table 1). *C. cinereum* δ^2^H_liver_ values were positively correlated with δ^2^H_month_ (*F*:4.6,DF = 37, *P* = 0.04) (Fig. 2). *S. magellanicus* δ^2^H_liver_ values were positively correlated with δ^2^H_month_ (*F*:11.6, DF = 16, *P* < 0.001), latitude (*F:*11.6, DF = 16, *P* = 0.01), and negatively correlated with elevation (*F:*11.6, DF = 16, *P* = 0.01) (Fig. 1). *T. aedon* δ^2^H_liver_ values were positively correlated with δ^2^H_month_ (*F:* 6.7, DF = 26, *P* < 0.001). *Z. capensis* δ^2^H_liver_ values were positively correlated with δ^2^H_month_ (*F:*16.1, DF= 27, *P* < 0.01) (Fig. 2), negatively correlated with elevation of capture (*F:*16.1, DF = 27, *P* < 0.001) (Fig. 1), and negatively correlated with latitude of capture (*F:*16.1, DF = 27, *P* < 0.01).

**Table 2.**
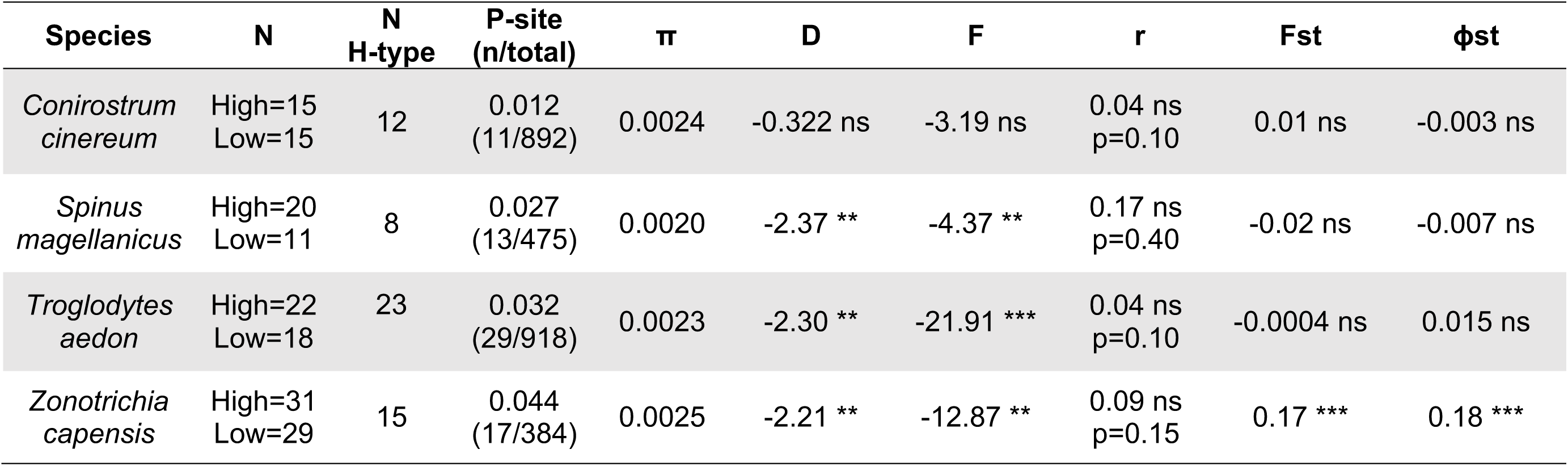
Summary of mtDNA analyses by species for low (<1000 m) and high (>3000 m) elevation specimens. Haplotypes (N H-Type), proportion of polymorphic sites (P-site), nucleotide diversity (p), Tajima’s D (D), Fu’s F (F), raggedness (r), Fst and ϕst are reported. Significant values accompanied by asterisks.

**Figure 2.**
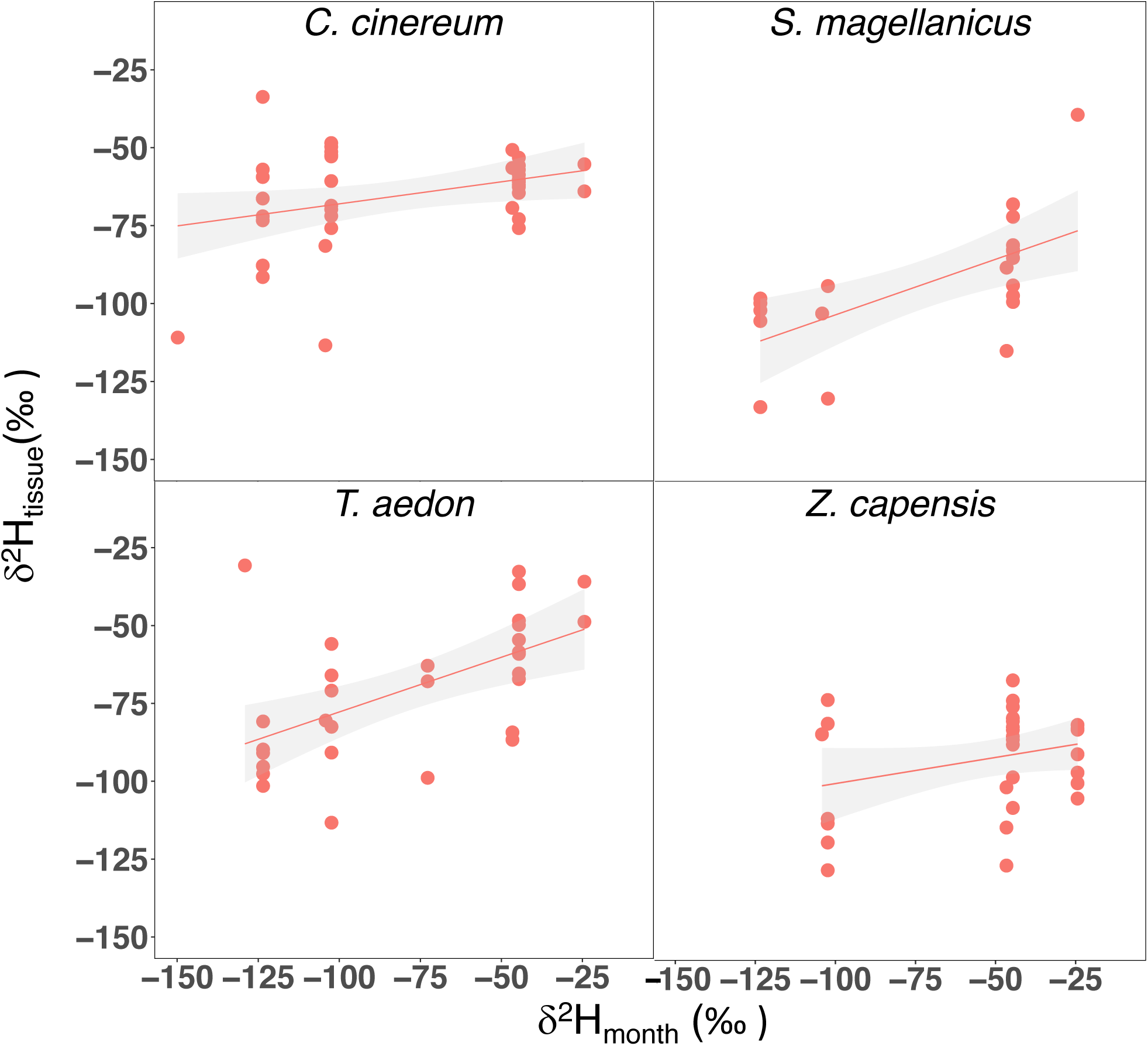
δ^2^H_liver_ plotted against 2006–2008 monthly mean precipitation δ^2^H (δ^2^H_month_) values from Marcapomacocha, Peru (IAEA/WMO, 2015); sample sizes for each species are reported in Table 2. Best-fit lines are shown for linear regressions that are significant at p < 0.05. Gray bands represent 95% confidence intervals.

Comparisons of δ^2^H values among tissue types are reported in Supplementary Table S2. Interspecific comparisons of δ^2^H_feather_ values are reported in supplementary Table S3.

### Morphometric comparisons

Mean morphometric measurements (culmen, tail, tarsus, and wing chord) are reported for low (< 1000 m) and high (> 3000 m) elevational bins for each species in Table S4. Wing length differed between elevations for *S. magellanicus* (*X^2^:*22.2, DF = 1, *P* = <0.001); tail length differed between elevations for *T. aedon* (*F*:6.7, DF = 1, *P* = 0.02) (Table S4). We found no measurement differences between elevational groups for *C. cinereum* or *Z. capensis*. The principal component analysis illustrates the overall findings of morphological differentiation in *S. magellanicus* and *T. aedon*, but not in the other two species (Fig. 3).

**Figure 3.**
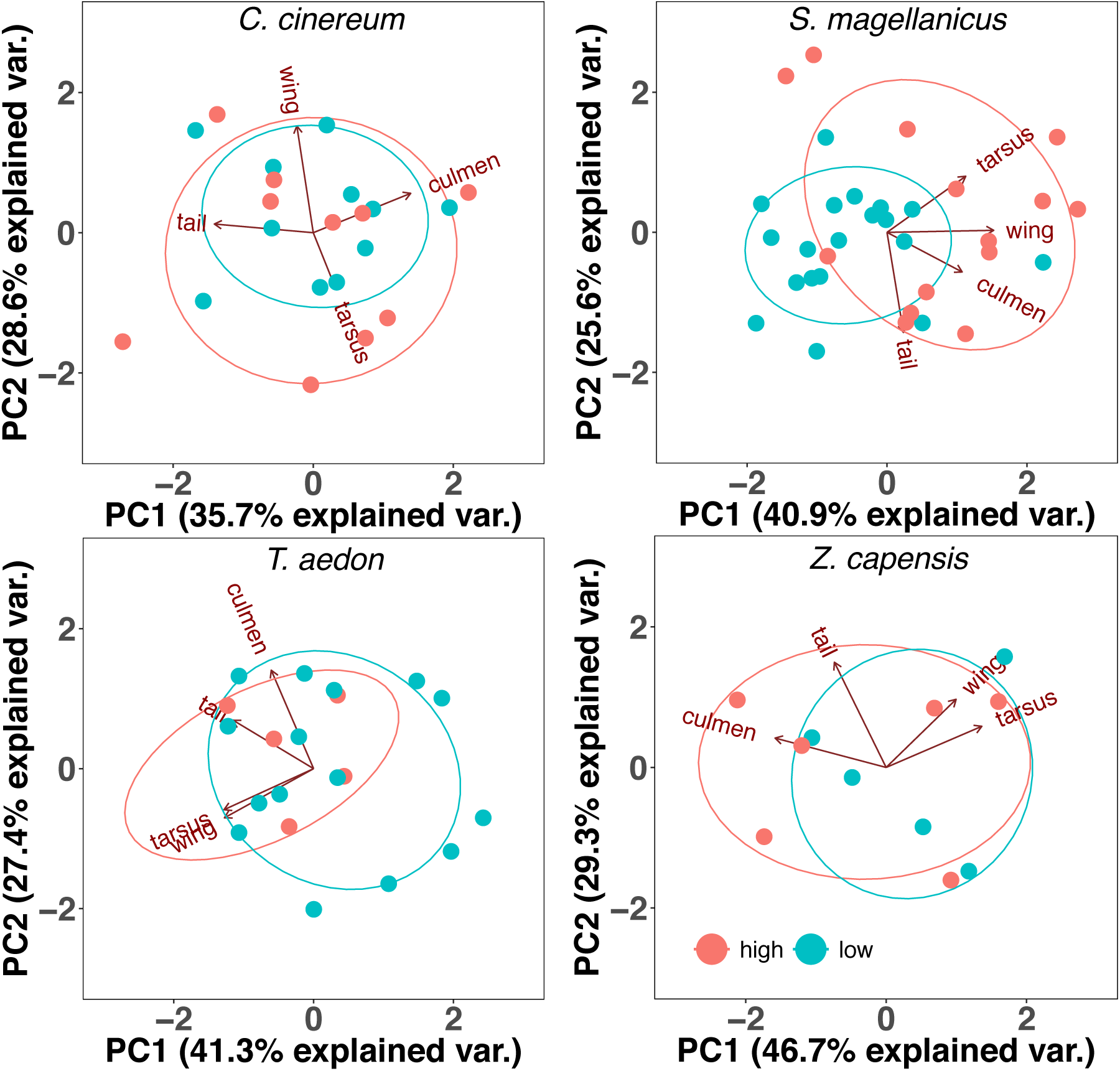
Principal component analyses of four morphological measurements: bill length, tarsus, wing chord, and tail length in millimeters for *C. cinereum* (n = 21), *S. magellanicus* (n = 34), *T. aedon* (n = 22), and *Z. capensis* (n = 11), grouped into (red) high elevation (>3000m) and (blue) low elevation (<1000m).

Hand-wing-index differed in all pairwise comparisons between species except those of *Z. capensis* with *C. cinereum* and *T. aedon*, respectively (Fig. S4). Those data indicate highest vagility in *S. magellanicus*, followed by *C. cinereum, Z. capensis*, and *T. aedon*. Species variation in flight muscle mass (corrected for body size) was consistent with the latter finding (Fig. S4). Flight muscle mass differed between species in all comparisons except the one between *Z. capensis* and *C. cinereum*.

### mtDNA structure and demography

Fst and Φst statistics were only significantly non-zero for the comparison between high (n = 31) and low elevation (n = 29) groups of *Z. capensis* (Table 2). Raggedness of mismatch distributions was not significant for any of the four species (Table 2, Fig. 4), which is consistent with the null hypothesis of recent demographic expansion. Tajima’s *D* and Fu’s *F* statistics were significantly negative, suggesting recent demographic expansion for *S. magellanicus, T. aedon*, and *Z. capensis*, but not for *C. cinereum* (Table 2, Fig. 4).

**Figure 4.**
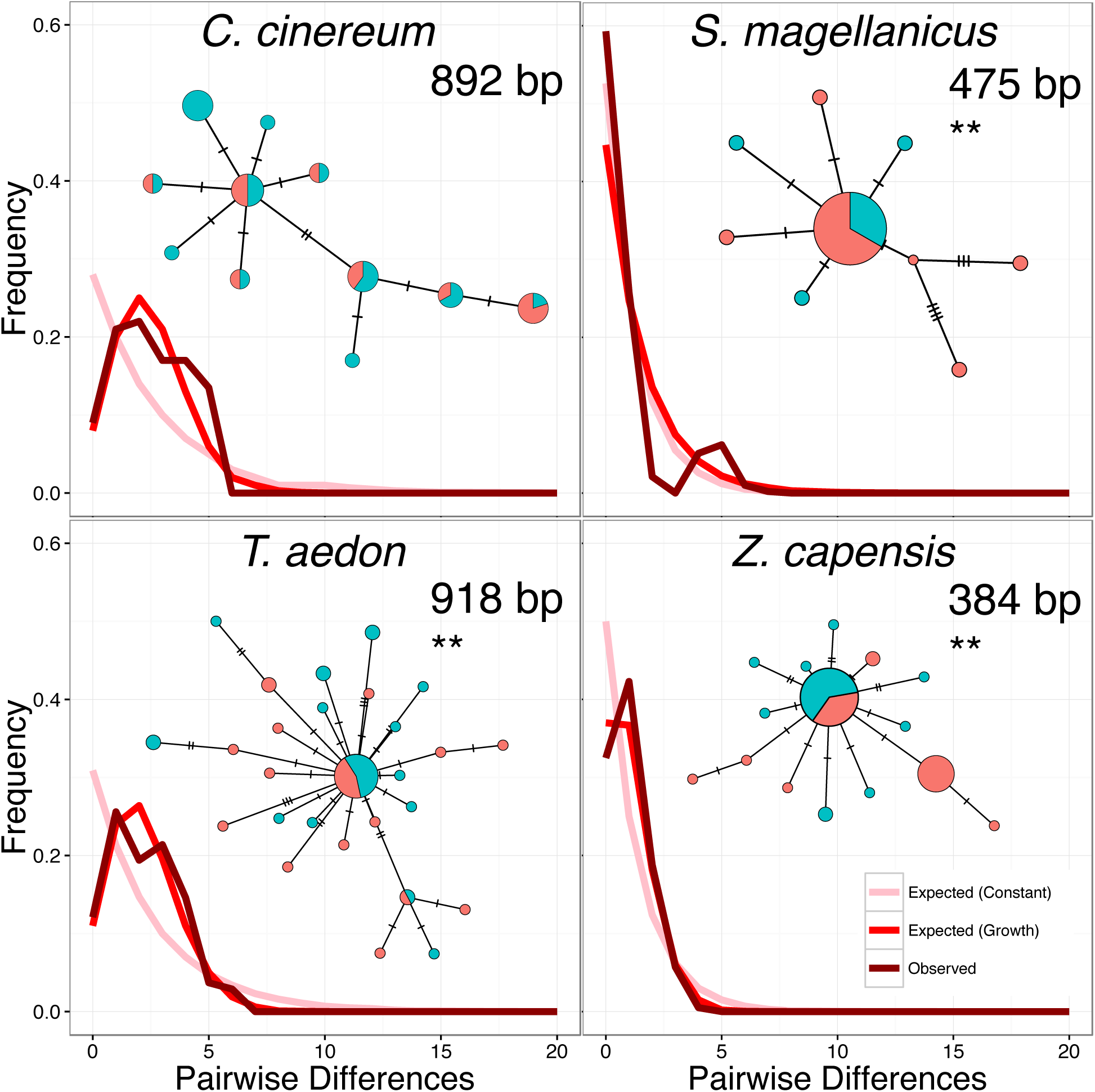
Mismatch distributions obtained from mtDNA loci (*ND2* or ND3) of high (>3000 m) and low (<1000 m) elevation specimens of *C. cinereum, S. magellanicus, T. aedon*, and *Z. capensis*. Haplotype networks colored by elevation group: high elevation (red), low elevation (blue) accompany each mismatch distribution. Double-asterisks indicate that both Tajima’s D and Fu’s F tests for population expansion, based on haplotype frequency spectra, were significant for that species.

## Discussion

Elevational movement, elevational genetic differentiation, and evidence of recent population expansion occur to varying degrees in our four study taxa, suggesting that each may play a role in causing exceptions to Janzen’s Rule. Tissue δ^2^H patterns associated with elevational movement were observed in two species (*C. cinereum* and *S. magellanicus*) that had no mtDNA population structure, only one of which (*S. magellanicus*) showed morphological differentiation. Two other species showed isotopic patterns associated with sedentary habits; one of those species exhibited mtDNA population structure (*Z. capensis*) while the other exhibited morphological differentiation (*T. aedon*). Three out of the four focal species showed signs of recent population expansion by all three indices tested. In the following sections, we examine tissue δ^2^H patterns associated with elevational movement and we consider how differing environmental processes might obscure or maintain these patterns. We delineate subcategories of elevational generalists (sedentary and migratory) to illustrate potential evolutionary consequences of short distance migration along environmental gradients.

### Tissue δ^2^H patterns

Variation in *T. aedon* and *Z. capensis* feather δ^2^H with elevation of capture (Fig.1) conforms with expected elevational patterns in precipitation δ^2^H, which suggests these species are generally sedentary. Both species are terrestrial foragers with morphological characteristics indicating low to moderate vagility (Table S1). For both species, the best linear models to explain variation in δ^2^H _contour_ and δ^2^H_secondary_ included only elevation as a predictor variable (Table 1). This indicates that individuals of these species had molted at or near the elevation of capture. It further suggests that the season of molt was consistent among individuals such that the elevational signal was not overwhelmed by seasonal fluctuations in precipitation δ^2^H. For the other two species, *S. magellanicus* and *C. cinereum*, elevation did not explain variation in feather δ^2^H, suggesting that individuals of those species underwent elevational movements between the time of molt and the time of sampling. A plausible alternative explanation would be that the season of molt is more variable among individuals of the latter two species, but there is no evidence for that in our molt data (Fig. S2), so we favor the conclusion that *S. magellanicus* and *C. cinereum* are elevational migrants.

Unlike feather δ^2^H values, δ^2^H_liver_ integrate seasonal variation in precipitation δ^2^H because this tissue is metabolically active and has a rapid isotopic incorporation rate, integrating ecological information over 1–2 weeks prior to capture for an endotherm the size of a songbird (Martínez del Rio et al. 2009, Wolf et al. 2009). δ^2^H_liver_ showed significant positive associations with δ^2^H_month_ for all species (Fig. 2; Table 1). Elevation of capture was included in the best models for δ^2^H_liver_ for all species except *C. cinereum* (Table 1), though elevation was significant only for *Z. capensis* (Fig. 1). It is possible that the lack of elevational trend in δ^2^H_liver_ for *C. cinereum* might have occurred due to elevational movements within the weeks before sampling, but the generally modest relationship between elevation and δ^2^H_liver_ may have other causes (see below).

Any seasonal variation in precipitation δ^2^H that was not captured by our temporal index (δ^2^H_month_) may have dampened the expected elevational trends in δ^2^H_liver_. δ^2^H_month_ provided an index of seasonal flux in precipitation δ^2^H that was derived from three years (2006-2008) of data at a single high elevation site, Marcapomacocha (∼4400 meters). It is possible that the Marcapomacocha data poorly represented seasonal fluctuations in precipitation δ^2^H at other elevations or in other years. Thus, future studies would greatly benefit from the collection of additional precipitation δ^2^H data along Andean elevation gradients. Additional problems with interpretation of our models could have occurred if sampling during the wet or dry season were concentrated at high or low elevation; however, we consider these potential sources of bias unlikely to have driven our results because our δ^2^H data came from specimens that were collected across the entire elevational gradient during both wet and dry periods for all species (Fig. 1, Fig. 2).

### Morphological and genetic differentiation

The division between sedentary and migratory modes of elevational generalism reflected in tissue δ^2^H patterns is likely mirrored in flight capabilities. Depending on foraging strategies and local ecologies, sedentary birds are predicted to be less vagile than their migratory counterparts, traits that should be reflected in the flight apparatus (flight muscle size and hand wing index; Fig. S4) and foraging stratum (Table S1). Larger flight muscle mass, higher hand wing index, and less terrestrial foraging ecology in *C. cinereum* and *S. magellanicus* relative to *T. aedon* and *Z. capensis* generally support this dichotomy, although there was some overlap in hand-wing index in *Z. capensis*–*T. aedon*/*C. cinereum* comparisons (Fig. S4) and flight muscle size between *C. cinereum* and *Z. capensis* (Table S1; Fig. S4).

Differences in sedentary and elevational-migratory habits should be further reflected in their respective levels of within-species population-genetic structure. Given enough time, we expect sedentary elevational generalists to have developed genetic structure between high- and low-elevation populations. Conversely, migratory behavior in elevational generalists should maintain or enhance gene flow, effectively washing out any incipient population structure. We expect that tests for population level differentiation within this two-mode framework will provide insight into the ecologies and evolutionary trajectories of bird species that are elevational generalists.

Our morphological tests showed significant differentiation between high and low elevation *T. aedon* in tail length (Fig. 3 and S5). A trend of larger appendages at higher elevations has been previously reported in another Andean bird, the Torrent Duck (Gutiérrez-Pinto et al. 2014). The morphometric disparity, in combination with our δ^2^H data, is in agreement with our hypothesis regarding the link between sedentary habit and elevational genetic differentiation. In contrast, *C. cinereum* exhibited no differentiation between high- and low-elevation specimens in the four characters we measured. This lack of differentiation is consistent with our predictions for an elevational generalist that is also an elevational migrant. *S. magellanicus* showed significant morphological differentiation in wing chord length between high- and low-elevation specimens. This morphological population structure conflicts with an isotopic pattern indicating elevational movement. The larger wing-chord sizes at high elevation could be the result of selection on wing-size that was strong enough to overcome gene flow (Smith et al. 2004, Gutiérrez-Pinto et al. 2014, Benham and Witt 2016). An alternative possibility is that the traits we measured exhibit high levels of phenotypic plasticity in response to elevation-specific pressures, but we consider this possibility to be unlikely. Phenotypically plastic traits could produce similar elevational patterns in the absence of genetic population structure by changes in gene expression alone (Przybylo et al. 2000, Cheviron et al. 2008), but there is evidence that morphometric traits remain highly heritable despite this possibility (Boag 1983, Keller et al. 2001).

Population structure can persist locally along contiguous elevational distributions, effectively selecting against unfit immigrants (Cheviron and Brumfield 2009, Cheviron et al. 2014). Yet, if average dispersal distances are large, these clines are unlikely to form. To test for genetic differentiation, we analyzed *ND2* or *ND3* mtDNA sequence data from all four species. Fst and Φst values confirmed previously reported population structure between high and low elevation *Z. capensis*, corroborating a sedentary habit for this species (Fig. 4) (Cheviron and Brumfield 2009). Fst and Φst values were not significant for *T. aedon*, which was somewhat surprising considering the sedentary lifestyle suggested by our isotopic and morphometric data. Analysis of β-hemoglobin gene variation in *T. aedon* across the same elevational transect studied here found substantial elevational population structure (Galen et al. 2015). mtDNA analyzed here was unstructured with respect to elevation, as were the vast majority of nuclear protein-coding genes analyzed by Galen et al. (2015). As in *T. aedon*, mtDNA sequence data from *C. cinereum* and the *S. magellanicus* showed no population structure between high- and low-elevation groups.

Signals of recent demographic expansions, as indicated by mismatch distributions, Tajima’s D, and Fu’s F test statistics (Table 2, Fig. 4) were present in three of the four focal species (*S. magellanicus, T. aedon,* and *Z. capensis*). These demographic expansions, if accompanied by expansions of the elevational range, potentially explain exceptions to Janzen’s Rule (1967). Considering the physical landscape of the western Andes, this expansion likely originated in high elevation environments that are diverse and productive relative to dry coastal zones that are depauperate and may have fewer competitors. Published phylogenies for tanagers (including *C. cinereum*) (Burns et al. 2014), siskins (including *S. magellanicus*) (Beckman and Witt 2015), South American *T. aedon* populations (Galen and Witt 2014, Galen et al. 2015) and *Z. capensis* populations (Lougheed et al. 2013) are all consistent with montane origins and subsequent, downslope range expansions in western Peru. As a caveat, it should be noted that false inference of population expansion from mtDNA haplotype frequency spectra can be caused by other demographic events, such as selective sweeps (Fay and Wu 2000, Wakeley and Aliacar 2001, Przeworski 2002). Moreover, the high prevalence of apparent range expansions among these elevational generalists contrasts with previous findings for Andean cloud-forest specialist species; haplotype frequency spectra consistent with population expansion were found in only a small fraction of subpopulations for *Thamnophilus caerulescens* (Brumfield 2005), *Metallura tyrianthina* (Benham et al. 2015, Benham and Witt 2016), and *Premnoplex brunnescens* (Valderrama et al. 2014). Two species of brush-finches (*Buarremon*) that are restricted to mid-elevations appear to have undergone recent expansions, but the evidence was considered to be equivocal (Cadena 2007).

Our mtDNA data provides insights into the timing of the inferred range expansions. Fossil-calibrated divergence rates such as the oft-used 2% per million years (Lovette 2004, Weir and Schluter 2008) are known to overestimate the ages of recent events (Arbogast et al. 2002, Ho et al. 2015). Therefore, we used a pedigree-based substitution rate (3.13 x 10^-7^ mutations/site/year) derived from chicken mtDNA genomes to estimate dates of population expansion (Alexander et al. 2015). Applying this rate to our mtDNA data, we estimated expansion to have occurred ∼3.5 Kya (*T. aedon*) to ∼34 Kya (*S. magellanicus*).

### Possible anthropological influence on expansion

Considering that our estimates of the timing of population expansion are as recent as ∼3.5 Kya, it is possible that these expansions may have coincided with human activity in this region. The lower west slope of the central Peruvian Andes is one of the driest places on Earth, with very limited natural bird habitats away from the immediate vicinity of rivers and streams sourced from the high Andes. Isolated patches of ‘lomas’ vegetation that depend on water from persistent coastal fog comprise one exception (Rundel and Dillon 1998). With the exception of lomas patches, the expansion of bird habitats away from rivers would have occurred only recently, following the implementation of sophisticated irrigation systems by the Paracas people, which also occurred ∼3.5 Kya (Hesse and Baade 2009). Regardless of whether it directly caused signals of expansion in our genetic data, the expansion of bird populations spurred by water diversion and irrigation on formerly arid land should be considered likely. Currently, *T. aedon* and *Z. capensis,* though widespread in undeveloped areas, are facultative human commensalists (Ruiz et al. 2002, Newhouse et al. 2008). Whether or not agricultural development facilitated expansion to lower and dryer portions of the western Andean slopes, our findings of recent expansion, individual movements, and ongoing diversification indicate that evolutionary instability is inherent to broad elevational ranges, at least for tropical songbird species.

## Acknowledgements

We thank the UNM Center for Stable Isotopes (Z. Sharp, V. Autedori), M. Baumann, P. Benham, L. Burkemper, R. Dickerman, S. DuBay, A. Johnson, J. Noble, J. A. Otero, C. G. Schmitt C. J. Schmitt, D. Schmitt, T. Valqui, and N. Wright. Funding included NSF DEB-1146491.

Supplementary material (Appendix EXXXXX at < www.oikosoffice.lu.se/appendix/). Appendix 1–8.

